# IL-4 receptor targeting as an effective immunotherapy against triple-negative breast cancer

**DOI:** 10.1101/2020.08.05.238824

**Authors:** Sadiya Parveen, Sumit Siddharth, Laurene S Cheung, Alok Kumar, John R Murphy, Dipali Sharma, William R Bishai

## Abstract

In many solid tumors including triple-negative breast cancer (TNBC), IL-4 receptor (IL-4R) upregulation has been shown to promote cancer cell proliferation, apoptotic resistance, metastatic potential and a Th2 response in the tumor microenvironment (TME). Immunosuppressive cells in the TME including myeloid-derived suppressor cells (MDSCs) and tumor-associated macrophages (TAMs) also express the IL4-R. We hypothesized that selective depletion of IL4-R bearing cells in TNBC may have dual cytotoxic and immunotherapeutic benefit. To selectively target IL-4R^+^ cells, we genetically constructed, expressed and purified DABIL-4, a fusion protein toxin consisting of the catalytic and translocation domains of diphtheria toxin fused to murine IL-4. We found that DABIL-4 has potent and specific cytotoxic activity against TNBC cells *in vitro*. In murine TNBC models, DABIL-4 significantly reduced tumor growth, splenomegaly and lung metastases, and this was associated with reductions in MDSC, TAM and regulatory T-cells (Tregs) populations with a concomitant increase in the proportion of IFNγ^+^ CD8 T-cells. The anti-tumor activity of DABIL-4 was absent in IL-4R KO mice directly implicating IL-4R directed killing as the mechanism of anti-tumor activity. Moreover, NanoString analysis of DABIL-4 treated TNBC tumors revealed marked decline in mRNA transcripts that promote tumorigenesis and metastasis. Our findings demonstrate that DABIL-4 is a potent targeted antitumor agent which depletes both IL-4R bearing tumor cells as well as immunosuppressive cell populations in the TME.

**STATEMENT OF SIGNIFICANCE:** In solid tumors like breast cancer, Interleukin-4 receptor (IL-4R) expression in the tumor microenvironment aids tumor growth and metastasis. IL-4R expression upon host immune cells further dampens antitumor immunity. In this study, we have genetically constructed a fusion protein toxin, DABIL-4, composed of the catalytic and translocation domains of diphtheria toxin and murine IL-4. DABIL-4 showed specific cytotoxicity against triple-negative breast cancer (TNBC) cells in vitro. DABIL-4 also markedly inhibited TNBC tumor growth and metastasis in vivo. The primary activity of DABIL-4 was found to be depletion of IL-4R+ immune cells in combination with direct elimination of tumor cells. In conclusion, DABIL-4 targeting of both tumor and immunosuppressive host cells is a versatile and effective treatment strategy for TNBC.

## INTRODUCTION

Interleukin 4 (IL-4) is an important pleiotropic cytokine primarily secreted by activated Th2 lymphocytes, basophils, mast cells, and eosinophils. IL-4 promotes differentiation of Th2 cells, upregulates the expression of major histocompatibility complexes and the IL-4 receptor, and regulates immunoglobulin class switching, especially IgE production, making it an important cytokine in allergic responses (1). IL-4 functions by binding to three different classes of IL-4 receptors (IL-4R). The Type I receptor is primarily found on T-cells, basophils, mast cells, NK cells, and most B cells and consists of the IL4Rα and common gamma C (γc) subunits. The Type II receptor, comprised of the IL-13Rα1 and IL-4Rα subunits, is expressed on tumor cells. And the Type III receptor present on B-cells and monocytes consists of the IL-4Rα, IL-13Rα1 and γc subunits (2).

Upregulation of the IL-4R (CD124) has been shown in multiple human and murine malignancies including glioma, lung, breast, pancreatic, bladder, colon and ovarian cancers (2). The binding of IL-4 to the IL-4R recruits and phosphorylates tyrosine kinases Jak1/2 and Tyk2, which then activate the PI3K/AKT, MAPK, and Jak/STAT6 signaling pathways (3). Upon activation, these pathways promote cancer cell proliferation, resistance to apoptosis, and metastatic potential (3, 4). In the TME, IL-4 elicits increased Th2 responses and activates myeloid-derived suppressor cells (MDSCs) and tumor-associated macrophages (TAMs)—both of which express the IL-4R--to promote immunosuppression and angiogenesis (2). Therapeutic targeting of the IL-4/IL-4R interaction and its consequences has led to effective antitumor strategies which include pharmacological inhibition of the downstream pathways (AKT, ERK, JAK/STAT6, and mTOR), monoclonal antibody blockade of IL-4 or the IL-4R; engineered IL-4R antagonists, and IL-4 fused with cytotoxic payloads (2, 3).

In breast cancer, up-regulation of IL-4/IL-4R signaling has been associated with poor prognosis in both humans and murine models (5). Additionally, IL-4 blockade has been shown to effectively down-regulate the mitogen-activated protein kinase (MAPK) pathway and reduce the growth and invasion of breast cancer cells (4, 6). Interestingly, TNBC cells were shown to secrete higher levels of IL-4 in the tumor milieu, compared to estrogen-receptor (ER)-positive breast cancer cells, contributing to their proliferation and metastatic potential (4), and recently an IL-4 mediated boost in glucose and glutamine metabolism was identified as a driver of TNBC cell growth (7). As TNBC tumors are difficult to treat owing to their inherent molecular heterogeneity and lack of targetable receptors (estrogen, progesterone and HER2) (8–10), we sought to interrupt the IL-4/IL4R signaling axis in TNBC using a toxin-based approach.

We recently reported the development of a second-generation version of denileukin diftitox (a diphtheria toxin-related IL-2 fusion protein) that significantly decreased the growth of melanoma tumors when tested as monotherapy or in sequential combination with checkpoint blockade (11). In the present study, we describe the construction, expression, purification and action of IL-4R targeted fusion protein toxin: s-DAB1-_389_-mIL-4 (referred to as DABIL-4 hereafter). DABIL-4 consists of the catalytic and translocation domain of diphtheria toxin contained in residues 1-389 genetically fused to murine IL-4 (mIL-4). We demonstrate that this fusion protein toxin specifically targets IL-4R^+^ murine TNBC cells *in vitro. In vivo* administration of DABIL-4 in murine 4T1 adenocarcinoma model showed significant reductions in tumor growth, splenomegaly and metastases to lung. We also observed a marked decline in the population of MDSCs, TAMs and Tregs with a concomitant increase in proportion of effector CD8^+^ T-cells in TME. In summary, we demonstrate that DABIL-4 is a potent cytotoxic agent against IL-4R^+^ cells that also depletes immunosuppressive cell populations in both the TME and spleen and thereby promotes anti-tumor immunity.

## RESULTS

### Genetic construction, expression, and purification of DABIL-4

For the genetic construction and expression of DABIL-4, we used a strategy recently developed in our laboratory to re-construct the structural gene encoding denileukin diftitox with the native diphtheria *tox* signal sequence and a mutated *tox* operator (11). Using a *Corynebacterium diphtheriae* expression system, we achieved constitutive expression of the fusion protein toxin and its secretion into culture medium as a fully folded, biologically active, monomeric recombinant protein (**Fig 1A**). We constructed a recombinant plasmid (pKN2.6Z-LC128) to express DABIL-4. This plasmid was generated by replacing the IL-2 coding sequence in pKN2.6Z-LC127 (11) with a synthetic gene encoding murine interleukin 4 (mIL-4) that was codon-optimized for *C. diphtheriae* within an *E. coli* / *C. diphtheriae* shuttle vector (**Fig. S1**). For expression and purification of the fusion protein toxin, pKN2.6Z-LC128 was transformed into the non-lysogenic, non-toxigenic *C. diphtheriae* strain C7s(-)^tox-^ strain. The fusion protein toxin, which is secreted into the medium with this expression system, was concentrated, adsorbed onto a Ni^2+^-NTA affinity column, eluted and further purified by gel permeation chromatography to achieve >98% purity (**Fig 1B**). The chimeric nature of the fusion toxin was confirmed by immunoblot with anti-DT, anti-mIL-4, and anti-His6 antibodies. As shown in **Fig 1C**, the 58 kDa band reacted with all the three antibodies confirming that the fusion protein toxin retained the expected antigenic determinants.

**Fig 1.**
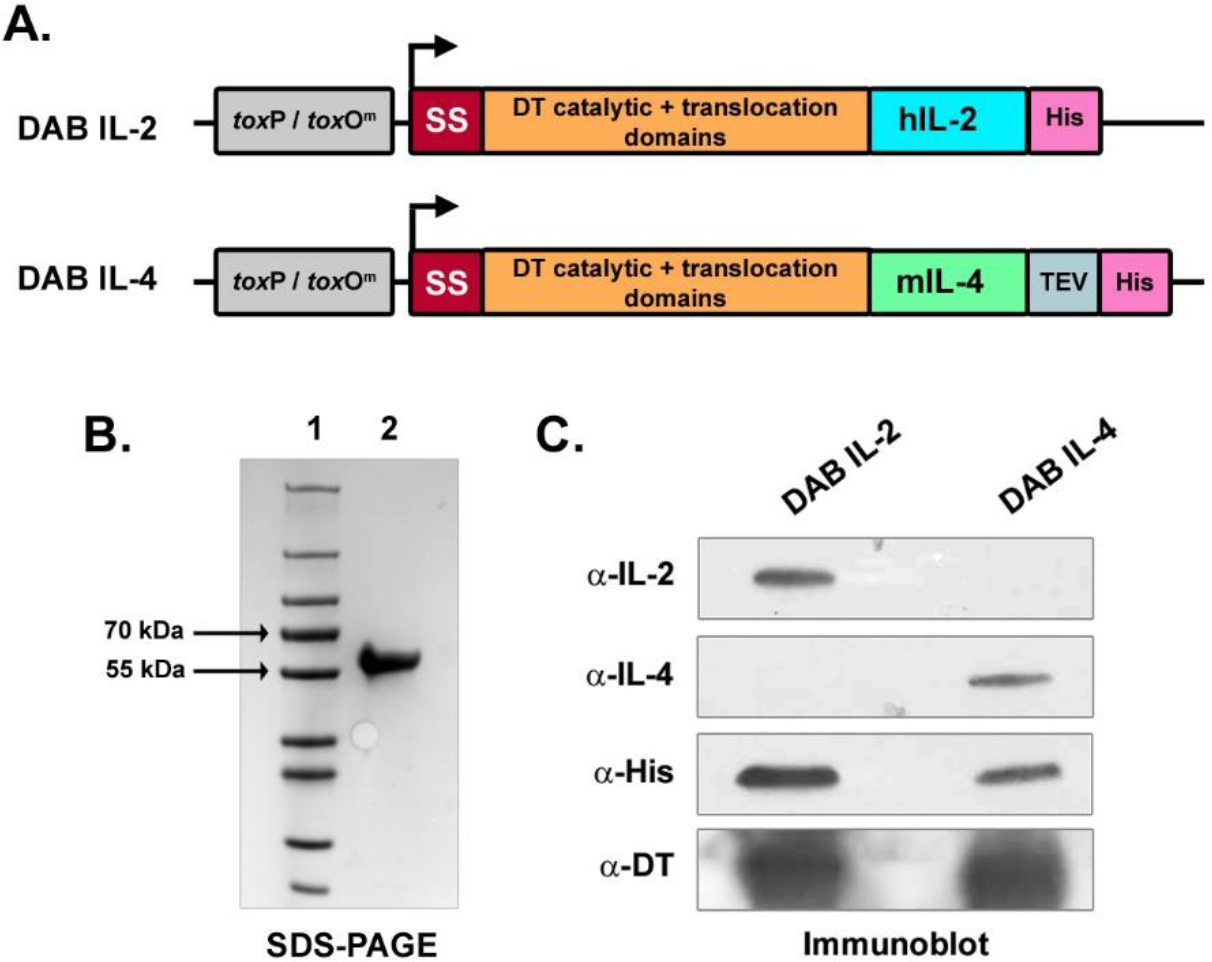
Genetic construction and purification of DABIL-4 fusion toxin using *C. diphtheriae*. **(A)** Schematic of DABIL-4 fusion toxin construct compared to DABIL-2 **(B)** Coomassie-stained SDS/PAGE gel after purification (lane 2). Lane 1 is molecular weight markers. Expected size of DABIL-4 is 58 kDa. **(C)** Immunoblot of the purified DABIL-2 and DABIL-4 fusion toxins. Staining with anti-DT antibody revealed diffuse bands for both DABIL-2 and DABIL-4 and arrow corresponds to the molecular weight of the fusion toxins.

### DABIL-4 selectively targets IL-4 receptor-expressing cancer cells in vitro

To evaluate whether DABIL-4 was cytotoxic for IL-4R-bearing cancer cells, we performed a dose-response analysis using 4T1 cells, a murine IL-4R^+^ triple-negative breast cancer cell line (6). Upon incubation with varying concentrations of DABIL-4 for 48 hours, 4T1 cells exhibited sensitivity to DABIL-4 in a dose-dependent manner with an IC_50_ of 2 nM (**Fig 2A**), while heat-inactivated DABIL-4 was completely non-toxic (**Fig 2B**). Another TNBC cell line, E0771 (12), also showed sensitivity to the fusion toxin albeit with a slightly higher IC_50_ value of 4 nM, a difference potentially attributable to different IL-4R expression levels and/or growth rates (**Fig S2**). 4T1 also exhibited sensitivity to DABIL-4 in a trypan blue dye exclusion assay with a calculated IC_50_ of 0.026 nM (**Fig 2C**). This discrepancy in IC_50_ values is likely due to the higher sensitivity of the dye exclusion assay compared to the colorimetric MTS assay. To demonstrate that the activity of DABIL-4 is mediated through the IL-4R, we examined the competitive inhibition of increasing concentrations of recombinant murine IL-4 upon the cytotoxicity of 25 nM DABIL-4. As shown in **Fig. 2D**, recombinant IL-4 completely blocked the cytotoxic activity, despite the presence of higher concentration of DABIL-4, demonstrating that the action of the fusion protein is mediated via the IL-4R/IL-4 interaction.

**Fig. 2.**
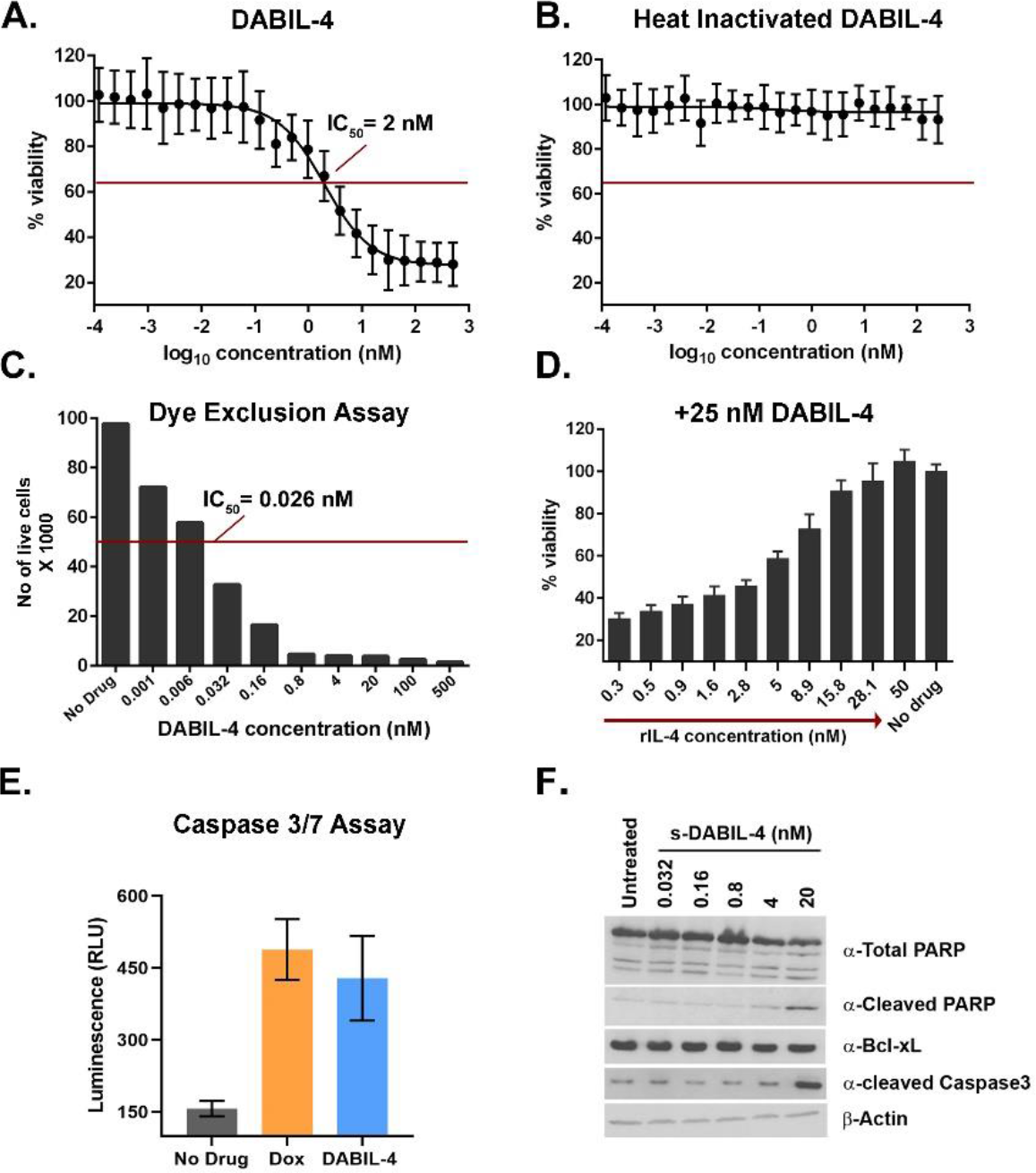
DABIL-4 exhibits cytotoxic activity against IL-4R^+^ 4T1 cells and induces apoptosis. **(A)** Activity of DABIL-4 in an MTS assay showed an IC_50_ of 2 nM while **(B)** post heat-inactivation, it showed no cytotoxicity. **(C)** DABIL-4 in trypan blue dye exclusion assay showed cytotoxicity with an IC_50_ of 0.026 nM. **(D)** Recombinant IL-4, in a dose-dependent fashion, blocked the cytotoxic activity of 25 nM DABIL-4 in an MTS assay. **(E)** Caspase 3/7 activity luminescence-based assay (Promega) performed with 4T1 cells treated with 20 nM DABIL-4. Caspase activity was comparable to doxorubicin, an agent known to cause apoptosis in 4T1 cells. **(F)** Immunoblot analysis of 4T1 cells treated with varying concentration of DABIL-4 showed upregulation of apoptotic markers (cleaved PARP and cleaved Caspase-3). β-actin was used as the loading control.

As previously reported, once internalized by the target cell, the diphtheria toxin catalytic domain ADP-ribosylates elongation factor 2 (EF-2) which leads to the inhibition of cellular protein synthesis, followed by induction of apoptosis and cell death within 48-72 hours (13). Using a chemiluminescence based assay, we demonstrated that DABIL-4 treatment led to 3-fold induction of caspase 3/7 activity in 4T1 cells which was comparable to the cytotoxic agent doxorubicin (Dox)(**Fig 2E**). Immunoblot analysis further demonstrated apoptosis induction in DABIL-4 treated cells revealing cleavage of both PARP and caspase3 (**Fig 2F**). These results confirm that DABIL-4 is a potent cytotoxic agent which induces apoptosis in the targeted cancer cells expressing the IL-4R.

### DABIL-4 inhibits tumor growth and prevents lung metastases in a murine TNBC model

The 4T1 mammary tumor model in syngeneic BALB/c mice shares many characteristic features with human breast cancer including progressive growth of primary tumors and the ability to metastasize to lungs, liver, bone and brain. Following orthotopic injection of as few as 10,000 cells into the mammary fat pads, palpable tumors appear within 7 days and metastases are observed by 18-28 days (14). We utilized the orthotopic 4T1 murine model to evaluate the cytotoxicity of DABIL-4 against 4T1 cells *in vivo* (**Fig 3A)**. Tumor bearing mice were treated by intraperitoneal injection on alternate days with either 10 μg DABIL-4 or PBS alone. As shown in **Figs. 3B** and **3C**, we observed significant reductions in tumor volume and weight in mice treated with DABIL-4. To rule out the strain specificity, we tested the efficacy of DABIL-4 in the orthotopic E0771 TNBC model generated in C57BL/6 strain background and observed that DABIL-4 significantly reduced E0771 tumor growth (**Fig S3**).

**Fig 3.**
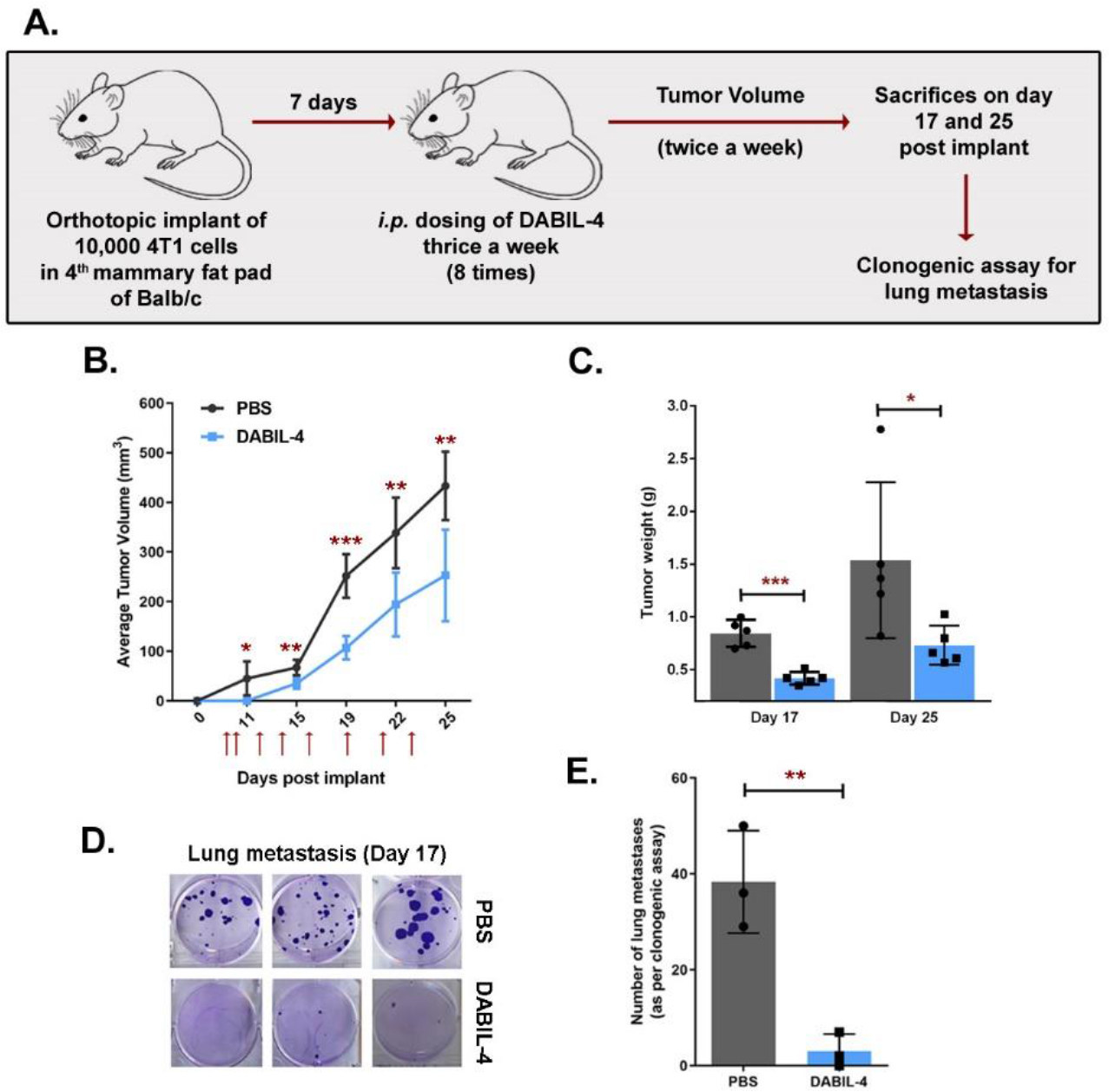
DABIL-4 shows anti-tumor activity in 4T1 adenocarcinoma model in WT background. **(A)** Schematic of the experiment. Treatment was given thrice weekly on days 7, 9, 12, 14, 16, 19, 21, 23 post tumor implantation. **(B)** DABIL-4 administration led to the decline in tumor volume (n=5), **(C)** tumor weight (n=5) and, **(D)** lung metastasis (n=3) both qualitatively and **(E)** quantitatively. Statistical significance for tumor growth and lung metastasis experiments was assessed by two-tailed unpaired student t-test considering an unequal distribution. Tumors were measured by electronic Vernier calipers. Red arrows indicate the days when treatment was administered. Data are shown as mean ± SD. *P < 0.05, **P < 0.01, ***P < 0.001.

4T1 tumors are also known to have high metastatic potential (14). To evaluate the impact of DABIL-4 upon metastatic progression, we harvested lungs from 4T1 tumor-bearing mice from both treated and untreated mice on day 17, isolated metastatic cancer cells and performed clonogenic assays. We observed a 12-fold reduction in the number of metastatic tumor colonies cultured from the lungs of DABIL-4 treated mice compared to the PBS-treated group (**Fig 3D** and **3E**). To further characterize the *in vivo* effect of DABIL-4 upon the metastatic potential, we analyzed total RNA isolated from the primary tumors using the NanoString PanCancer Immune Profiling Panel Assay. We noted significant declines in the level of several transcripts associated with metastasis (Cd36 (15), Egr3 (16-18) and Maf (19)) and tumorigenesis (Ccr2 (20), Ccr6 (21)) in many solid tumors including breast cancer (**Fig 4** and **Table 1**). We also observed an increased levels of the C200r1 transcript which is associated with inhibition of metastasis and tumorigenesis (22, 23)(**Fig 4** and **Table 1**). These observations indicate that DABIL-4 not only prevents primary tumor growth but also inhibits metastatic dissemination of 4T1 cells to secondary sites.

**Fig 4.**
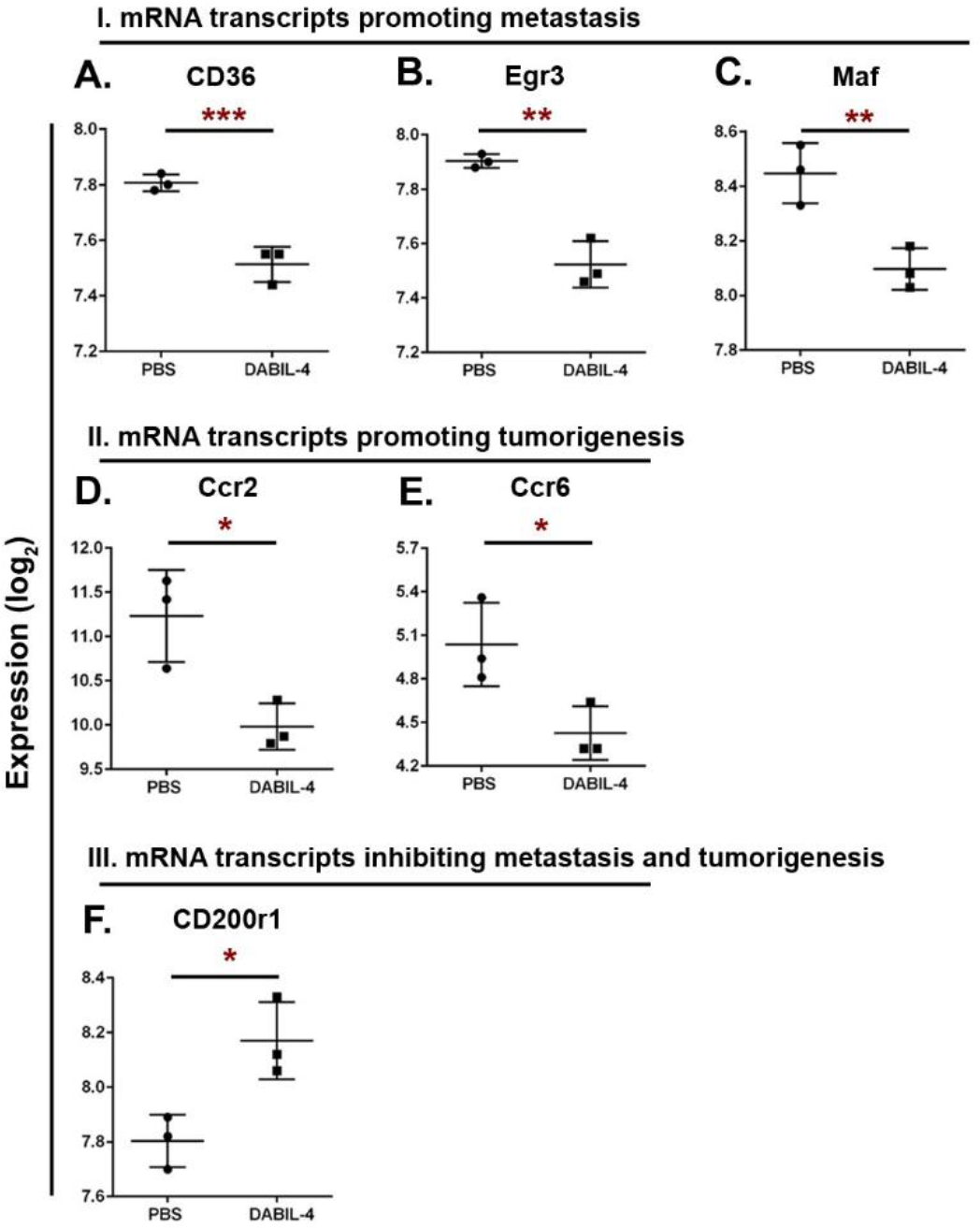
DABIL-4 monotherapy modulates level of mRNA transcripts associated with metastasis and/or tumorigenesis in tumor microenvironment. Total RNA isolated from 4T1 tumors (n=3) from both PBS- and DABIL-4 treated mice on day 17 and were subjected to NanoString PanCancer Immune Profiling Panel analysis and solver software. Expression profiles of transcripts promoting metastasis **(A-C)** or tumorigenesis **(D, E)** and transcript inhibiting metastasis and tumorigenesis **(F)** are shown for both treatment groups. RNA expression values are given as log2 and data are shown as mean ± SD. Statistically significant differences were defined by two-tailed unpaired student t-test considering an unequal distribution. *P < 0.05, **P < 0.01, ***P < 0.001.

**Table 1:** List of mRNA transcripts identified using NanoString PanCancer Immune Profiling Panel

| mRNA Transcript | Expression (log<sub>2</sub>) |  | P-Value | Pathway | Reference |
| --- | --- | --- | --- | --- | --- |
|  | PBS | s-DABIL-4 |  |  |  |
| **Cd36** | 7.81 | 7.51 | 0.005 | M | (15) |
| **Egr3** | 7.90 | 7.52 | 0.01 | M | (43) |
| **Maf** | 8.45 | 8.10 | 0.01 | M | (19) |
| **Ccr2** | 11.23 | 9.98 | 0.04 | T | (20) |
| **Ccr6** | 5.04 | 4.43 | 0.04 | T | (21) |
| **Cd200r1** | 7.80 | 8.17 | 0.03 | T/M | (22, 44) |
| **Gzmk** | 5.06 | 6.42 | 0.04 | I | (45) |
| **Il10ra** | 8.85 | 8.18 | 0.04 | I | (46) |

### DABIL-4 depletes MDSCs and TAMs in vivo thereby reducing immunosuppression in the TME

One of the pathologic hallmarks of the 4T1 model is splenomegaly (up to 8-fold increases in spleen weight) mostly due to granulocytic hyperplasia (24). We found that DABIL-4 treatment led to a 20% reduction in the spleen weights in 4T1 bearing mice suggesting that this fusion protein toxin also affects granulocytosis and tumor-associated leukemoid reactions in addition to tumor growth inhibition (**Fig 5A**). It is well known that IL-4R is not only expressed on 4T1 tumor cells but also upon immune cells, specifically myeloid-derived suppressor cells (MDSCs) and tumor-associated macrophages (TAMs). Both of these immune cell populations contribute to immunosuppression within the TME as well as metastasis (25–28). To investigate whether the observed inhibition of tumor growth was due to direct targeting of IL-4R^+^ tumor cells, to depletion of IL-4R^+^ MDSCs and TAMs, or to a combination of the two, we orthotopically implanted IL-4R^+^ 4T1 cells in an IL-4R knockout mouse strain (IL4Rα^-/-^) and treated with either PBS or DABIL-4 thrice weekly starting on day 7. We observed a delayed progression of 4T1 tumors in the IL-4R knockout mice. Moreover, treatment of tumor-bearing IL-4R knockout mice with DABIL-4 reduced spleen weights by 9% (vs 20% in WT) and tumor weights by 18% (vs 52% in WT), reductions that were less impressive than those we observed in wild type mice (**Fig 5B, 5C** and **5D**). These results in IL-4R knockout mice indicate that direct 4T1 cell killing by DABIL-4 contributes modestly to delayed tumor expansion. Further, they suggest that depletion of IL-4R^+^ immunosuppressive host cell populations is likely a key second mechanism of DABIL-4’s antitumor efficacy.

**Fig 5.**
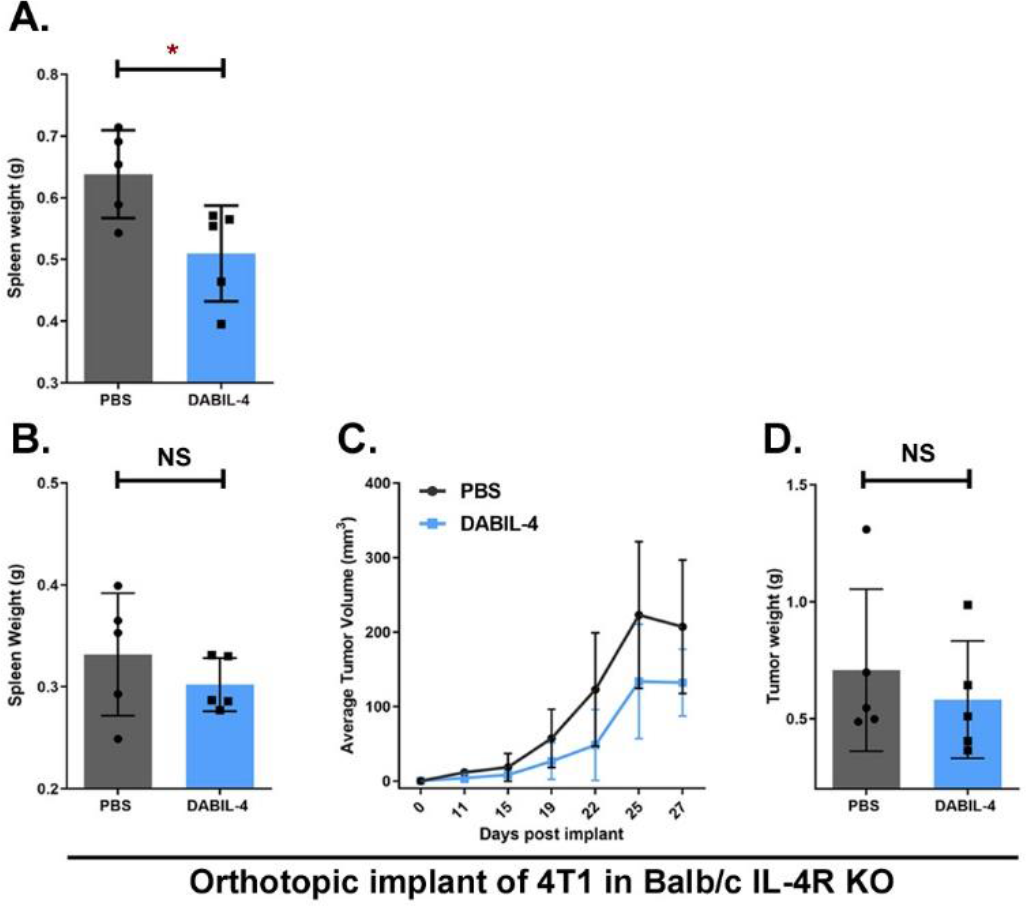
DABIL-4 shows no anti-tumor activity against 4T1 adenocarcinoma model in IL-4R knockout mice. **(A)** DABIL-4 administration in 4T1 bearing Balb/c mice showed reduction in spleen weights. In IL-4R KO mice, 10,000 4T1 cells were orthotopically implanted and allowed to establish. Starting day 7, DABIL-4 was given via i.p. thrice a week on alternate days for total of 8 doses. Mice were sacrificed on day 27. The fusion toxin administration did not significantly reduce **(B)** spleen weights, **(C)** tumor volumes and, **(D)** tumor weights. Statistical significance for these experiments was assessed by two-tailed unpaired student t-test considering an unequal distribution. Tumors were measured by electronic Vernier calipers. Data are shown as mean ± SD. *P < 0.05, NS=non-significant. N=5 mice per group.

To directly address the effects of DABIL-4 on host immune cell populations, we analyzed tumor/spleen from 4T1 tumor-bearing wild type mice using multicolor flow cytometry. As in **Fig. 3A,** mice began thrice weekly DABIL-4 or PBS treatment on day 7 after 4T1 tumor implantation. In spleen homogenates derived from DABIL-4 treated mice, we noted a 26% reduction in total myeloid cell population (CD11b^+^) (**Fig 6A**). This observation prompted us to investigate the drug’s impact upon two major myeloid cell subsets, MDSCs and macrophages. We first considered MDSCs and observed a marked decline in the total MDSC population (**Fig S4A**). Additionally, evaluation of two distinct MDSCs subsets; polymorphonuclear MDSCs (PMN-MDSCs; CD11b^+^ Ly6G^+^ Ly6C^low^ CD124^+^) and monocytic-MDSCs (M-MDSCs; CD11b^+^ Ly6G^-^ Ly6C^high^ CD124^+^) revealed a 60% reduction in IL-4R^+^ PMN-MDSCs population in DABIL-4 treated mice (**Fig 6B**), while the level of IL-4R^+^ M-MDSCs remained the same (**Fig S4B**). We also noted a sharp decline in the population of PMN-MDSCs subset expressing IL-10 (**Fig 6C**). This cytokine is known to skew anti-tumor T-cell responses towards a Th2 phenotype promoting tumor growth (29). Second, we evaluated macrophages and observed a marked increase in the overall population of macrophages (CD11b^+^ F4/80^+^) in spleen homogenates (**Fig S4C**). Upon further investigation, we found that DABIL-4 treatment had opposite effects upon M1- and M2-like macrophages. While we observed a significant decline in the population of M2-like macrophages (IL-4R^+^ CD206^+^) (**Fig 6D**), the population of pro-inflammatory CD86^+^ and TNFα^+^ M1-like macrophages increased (**Fig 6E** and **6F**).

**Fig 6.**
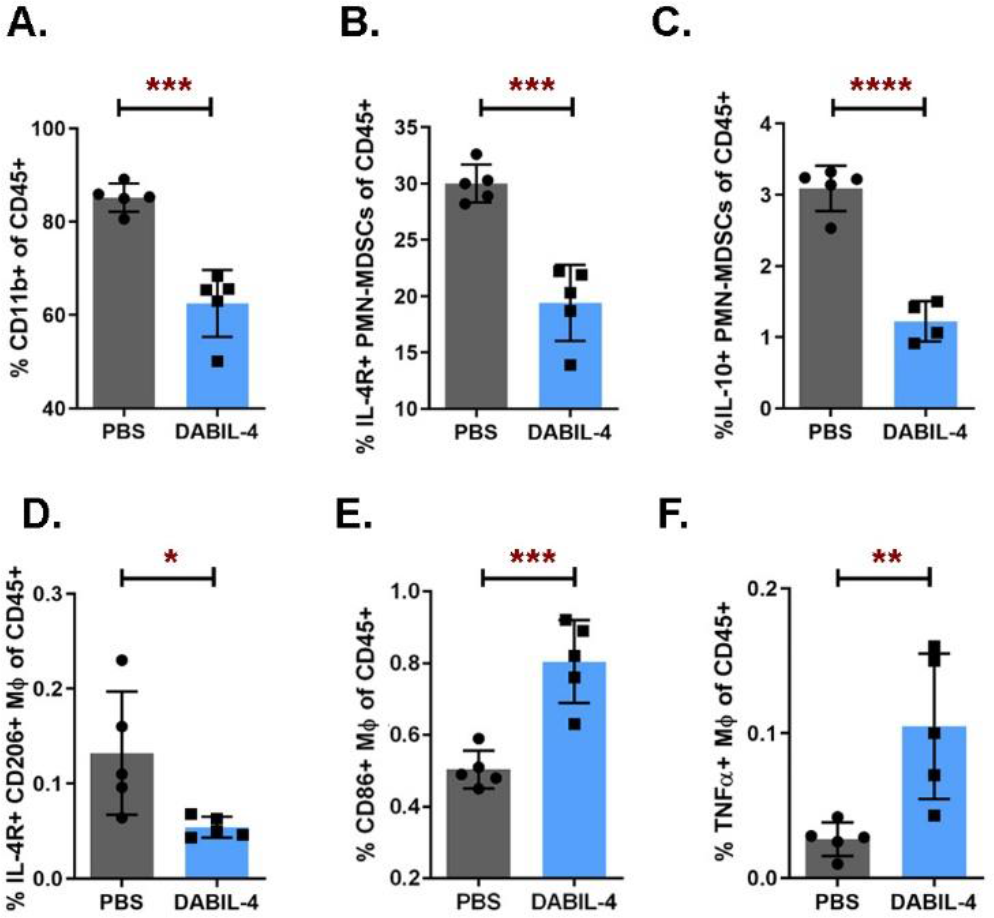
DABIL-4 administration depletes immunosuppressive myeloid cell population in spleen. As in **Fig 3A**, mice were treated with DABIL-4 thrice weekly beginning on day 7, and they were sacrificed on day 25. Single cell suspensions of spleens from both treatment groups were stained with specific antibodies and analyzed by flow cytometry (n=5). We found differences in the population of **(A)** CD11b^+^ cells, **(B)** IL-4R^+^ PMN-MDSCs, **(C)** IL-10^+^ PMN-MDSCs, **(D)** IL-4R^+^ CD206^+^ M2 macrophages, and **(E)** CD86^+^ M1 macrophages and **(F)** TNFα^+^ macrophages. Data are represented as mean ± SD and as percentage of CD45^+^ population (except panel D which is represented as “percentage population of macrophages”). Spleen were analyzed on day 26 post tumor implant. Statistical significance between the groups was assessed by two-tailed unpaired student t-test considering an unequal distribution. *P < 0.05, **P < 0.01, ***P < 0.001, ****P < 0.0001.

Similarly, in tumor homogenates, we noted reduction in IL-4R and CD206 expressing macrophages (**Fig S4D**) and an enrichment of TNFα^+^ macrophages (**Fig S4E**). These results suggest that DABIL-4 targets and depletes IL-4R^+^ MDSCs and macrophages thereby reducing their ability to suppress host mounted immune responses, resulting in a more robust anti-tumor response.

### DABIL-4 increases effector T cell population and their cytotoxic potential in vivo

Next, we investigated the effect of DABIL-4 administration upon both CD4^+^ and CD8^+^ lymphocytes in 4T1 bearing mice. As in **Fig. 3A,** mice began thrice weekly DABIL-4 or PBS treatment on day 7 after 4T1 tumor implantation and homogenates from spleens and tumors were analyzed using multicolor flow cytometry. In spleens, we detected enrichment of both CD4^+^ and CD8^+^ lymphocytes in DABIL-4 treated mice (**Fig S5A** and **S5B**). Both subsets also exhibited reduced expression of CD39 (**Fig S5C** and **S5D**), a phenotype associated with improved anti-tumor activity (30).

In the tumors, despite no apparent changes in total CD4^+^ and CD8^+^ T cells, we found a significant increase in CD44 and IFNγ expressing CD8^+^ lymphocytes in DABIL-4 treated mice (**Fig 7A** and **7B**). We also detected increased IFNγ and decreased CD39 expressing CD4^+^ T cells. (**Fig S6A** and **S6B**). There was also a significant decrease in the population of activated Treg cells (**Fig 7C**). These changes in lymphocyte subsets indicate that DABIL-4 treatment mice elicit enhanced anti-tumor responses in the TME. Accordingly, we detected significantly elevated levels of GzmK mRNA transcript in total RNAs isolated from these tumors (**Fig 7D** and **Table 1**) which may indicate either enhanced cytotoxic potential of lymphocytes or an overall increase in GzmK expressing lymphocytes population in the TME. The same analysis also revealed decline in IL-10 receptor expression in tumors from DABIL-4 treated mice (**Fig 7E** and **Table 1**).

**Fig 7.**
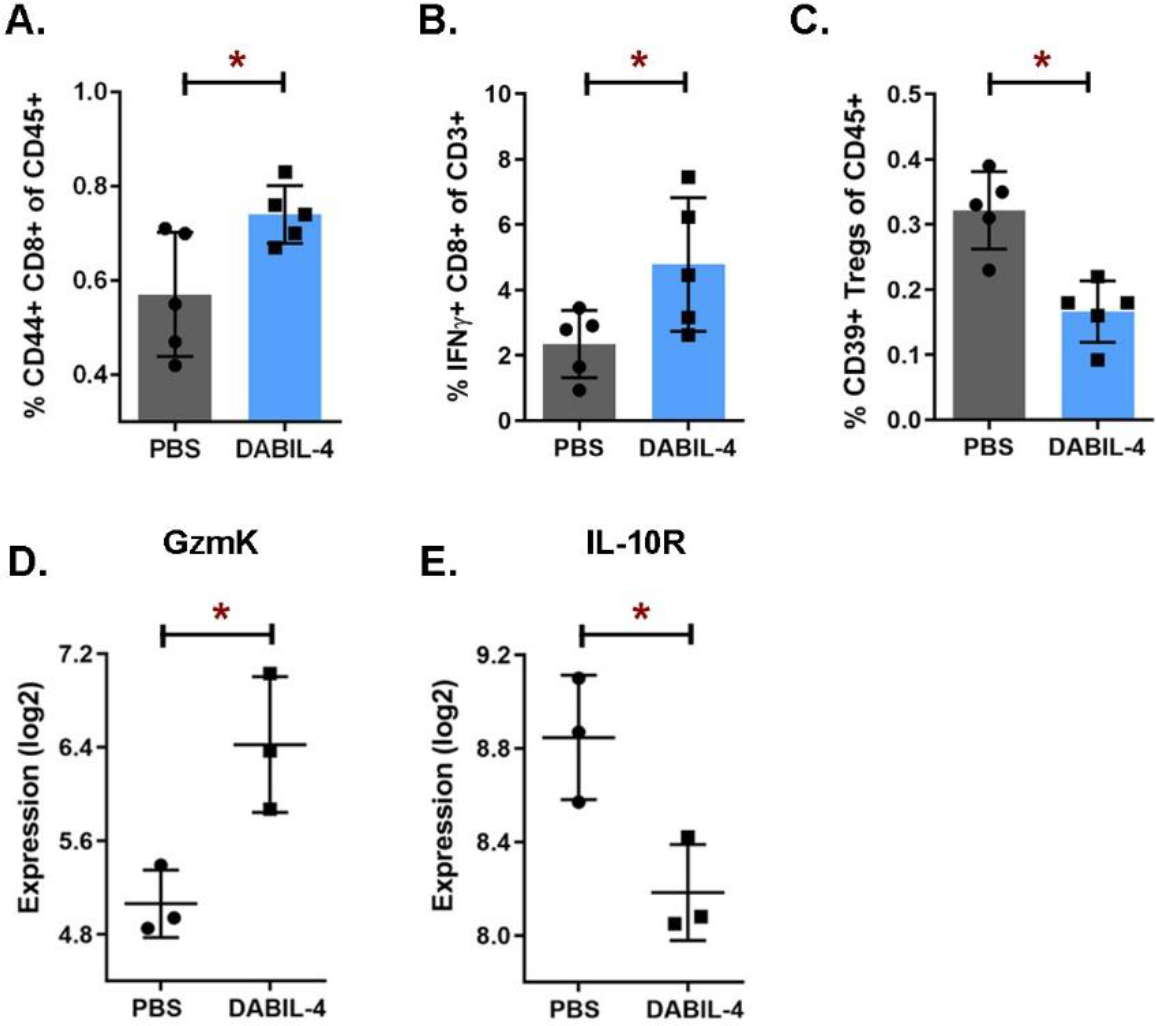
DABIL-4 treatment promotes infiltration of effector cell populations in tumor microenvironment, mediating anti-tumor response. As in Fig 3A, mice were treated with DABIL-4 thrice weekly beginning on day 7, and they were sacrificed on day 17 and 25. Single cell suspensions of the tumors isolated from both the groups were stained and were analyzed by flow cytometry (n=5). We found differences in the population of (A) activated CD44^+^ CD8^+^ T cells of CD45^+^, (B) IFNY^+^ CD8^+^ T cells of CD3^+^, and (C) CD39^+^ Tregs of CD45^+^ cells. Total RNA isolated from these tumors were subjected to NanoString PanCancer Immune Profiling Panel analysis and nSolver software (n=3). Expression profiles of (D) Granzyme-K and, (E) IL-10 receptor transcripts are shown for both treatment groups. RNA expression values are given as log2. All panels except panel B correspond to day 17 post tumor implantation while panel B corresponds to day 25. All data are shown as mean ± SD. Statistical significance between the groups was assessed by two-tailed unpaired student t-test considering an unequal distribution. *P < 0.05, **P < 0.01.

We also found an increase in frequency of an unconventional lymphocyte population, double-positive (DP) CD4^+^ CD8^+^ T-cells, in tumors isolated from DABIL-4 treated mice (p = 0.07) (**Fig S6C**). There was, however, a statistically significant increase in CD44 expressing DP lymphocytes (**Fig S6D**). Compared to CD4 and CD8 mono-positive T cells, DP lymphocytes exhibit characteristics similar to memory T lymphocytes, have a higher capacity to produce cytokines, and improved lytic potential (31, 32). Based on these observations we infer that DABIL-4 mediated depletion of immunosuppressive cell populations facilitates significant increases in the population of activated effector T cells in the TME.

## DISCUSSION

Here, we report the genetic construction, expression and purification of a diphtheria toxin-based fusion protein, DABIL-4, which targets the cell killing capability of diphtheria toxin to IL-4R-bearing cells. Using a *C. diphtheriae-based* expression host described earlier by Cheung et al. in 2019 (11), we expressed/purified DABIL-4 and demonstrated that the fusion toxin exhibits potent and specific cytotoxicity against IL-4R bearing TNBC cells *in vitro* and *in vivo*. The fusion toxin reduced tumor growth and lung metastases in a murine syngeneic TNBC adenocarcinoma model primarily by depleting IL-4R^+^ immune cell populations such as MDSCs thereby facilitating a robust anti-tumor T cell response, but it also showed modest depletion of the 4T1 tumor cells themselves.

Lakkis et al. reported a similar IL-4R^+^ diphtheria toxin-based fusion protein toxin from an *Escherichia coli-based* system and demonstrated targeted cytotoxicity to IL-4R bearing eukaryotic cells *in vitro* and a marked reduction in delayed type hypersensitivity *in vivo* (33). However, the therapeutic efficacy of the Lakkis et al. protein was not evaluated in preclinical animal tumor models and hence its effect upon host immune cells remained unknown. A similar immunotoxin, IL-4(38-37)-PE38KDEL, comprised of a circularly permutated human IL-4 fused with *Pseudomonas aeruginosa* exotoxin A, was found to be cytotoxic against the human breast cancer cell line, MDA-MB231, both *in vitro* and murine xenograft models (34). It was also tested in pre-clinical models of various human malignancies including breast cancer (2, 34). These studies with IL-4(38-37)-PE38KDEL were carried out in immunocompromised mice without an assessment of its effect on IL-4R^+^ immune cells, and thus its antitumor efficacy was largely attributed to direct killing of IL-4R-bearing tumor cells (34). In the present study, we demonstrated that DABIL-4 has a potent and direct cytotoxic activity against IL-4R^+^ positive TNBC cells *in vitro*. To assess the degree of direct tumor cell killing by DABIL-4, we tested it in IL-4R KO mice and found only modest reductions in tumor progression (Fig 5B-5D) suggesting the IL-4R^+^ immune cell depletion accounts for a significant degree of its antitumor efficacy. Indeed, we further demonstrated significant DABIL-4-mediated depletion of MDSCs and TAMs with concomitant increases in effector T cell populations and reductions in Tregs.

In the present study, we observed a selective depletion of splenic PMN-MDSCs while the population of splenic M-MDSCs remained the same in DABIL-4 treated mice. Interestingly, in spleen of 4T1 tumor bearing mice, PMN-MDSCs constituted the major MDSCs subset and exhibited higher proliferation rate compared to M-MDSCs (35, 36). This higher proliferation rate could be responsible for the increased susceptibility of PMN-MDSCs to DABIL-4 which inhibits translation, the major machinery required by the rapidly growing cells (37). In many cancers, M-MDSCs are shown to be abundant in the TME and tend to rapidly differentiate into TAMs (35). This dynamic conversion could be one of the reasons why we detected depletion of TAMs but not M-MDSCs in tumor. However, considering our limited ability to distinguish M-MDSCs from macrophages (38), the observed depletion of TAMs may also arise from targeting of both TAMs and M-MDSCs. Further investigation will be required to reach to a conclusion.

In 4T1 tumor bearing mice, the spleen acts as the primary site of MDSCs proliferation and sequestration (24, 36) from where these cells are then recruited to the TME in response to tumor-secreted chemokines (39). Indeed, evidences suggests that MDSCs-mediated suppression of T cells promote immunosuppression in the TME (40) (22). Accordingly, splenectomy has been shown to reduce the amount of tumor-infiltrating MDSC and to cause tumor regression in murine cancer models (41) suggesting that splenic MDSCs may also contribute to tumorigenesis (42), which explains why DABIL-4-mediated depletion of splenic MDSCs caused a robust anti-tumor response in the present study.

In this study, we demonstrated therapeutic efficacy of DABIL-4 as a monotherapy in murine model of triple-negative breast cancer. We propose that the therapeutic efficacy of DABIL-4 may be further potentiated by combining it with other effective immunotherapeutic approaches such as checkpoint inhibitors. While our study focused on breast cancer, there are several reasons to anticipate that DABIL-4 may offer a valuable immunotherapeutic approach against other solid tumors. First, we observed DABIL-4 mediated anti-tumor activity in both PDL1-negative (4T1) and positive (E0771) tumors models. Second, DABIL-4 showed efficacy even against well-established 4T1 murine tumors. The fusion toxin was active when therapy began either on day 7 (**Fig 3B**) or day 11 (**Fig S3**) post tumor implant. Third, IL-4R upregulation on immune and tumor cells and its role in tumorigenesis and metastasis is well-established in case of multiple solid tumors (2, 3), and DABIL-4 targets these tumor adaptations. In summary, DABIL-4 is a promising therapeutic that demonstrates both direct killing of IL-4R-positive tumor cells combined with depletion of IL-4R-bearing immunosuppressive cells such as MDSCs and TAMs. As such, It holds potential as dual cytotoxic and immunotherapeutic agent for the treatment of multiple cancers including triple-negative breast cancer and warrants further study.

## AUTHOR CONTRIBUTIONS

SP, JRM, DS and WRB conceptualized the study and designed the research approach; SP, SS, AK performed research; LC, JRM contributed new reagents/analytic tools; SP, JRM, DS, WRB analyzed data; and SP, JRM, DS, WRB wrote the paper, SP, SS, LC, AK, JRM, DS, WRB critically reviewed the manuscript.

## ACKNOWLEDGEMENTS

We gratefully acknowledge the support of NIH AI 130595, Tedco grants 2016-MII-3464 and 2019-MII-0518, the Abell Foundation and the Maryland Cigarette Restitution fund. We thank Dr. Geetha Srikrishna for editorial assistance.

## SUPPLEMENTRY FIGURES

**Fig S1.**
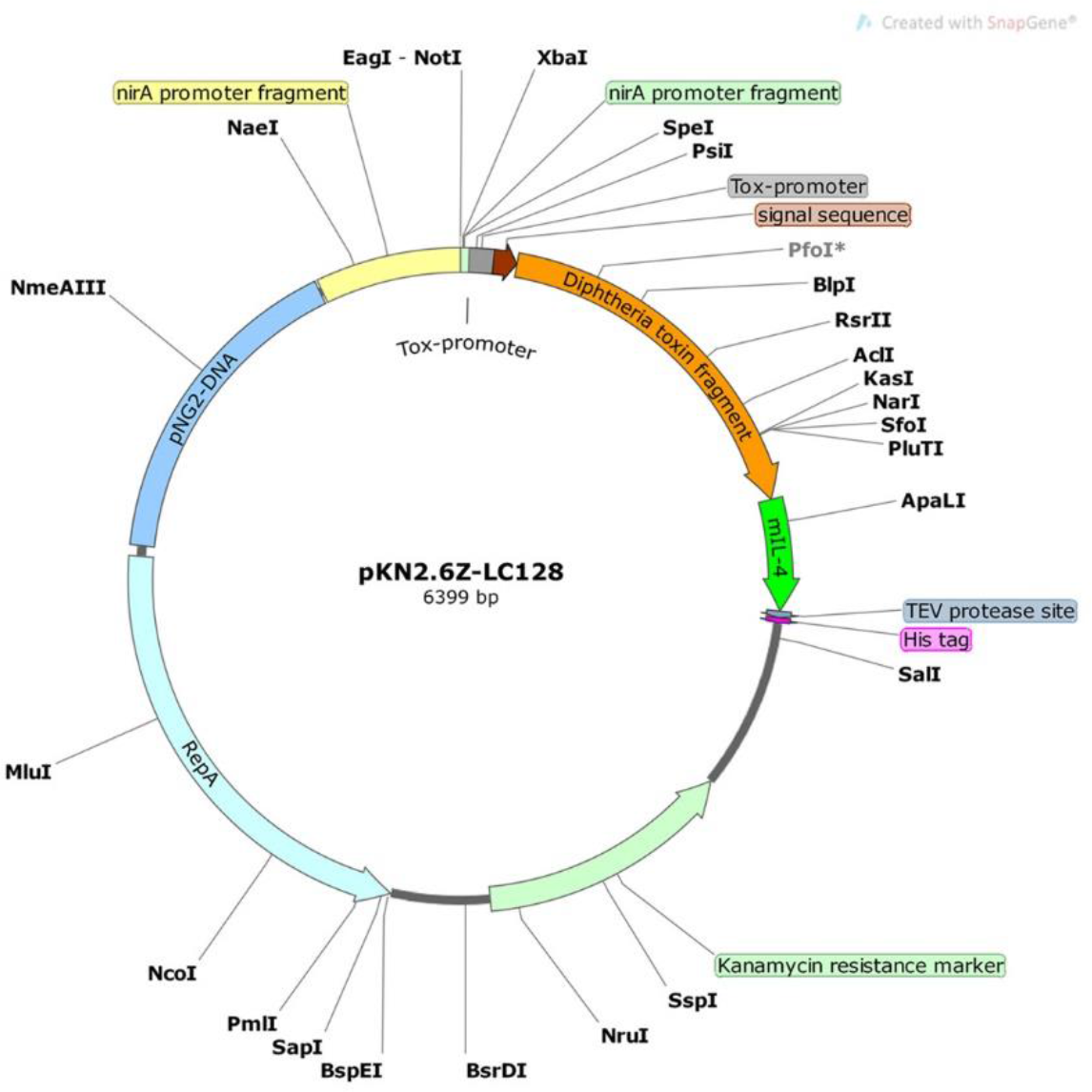
pKN2.6Z-LC128 shuttle vector plasmid map.

**Fig S2.**
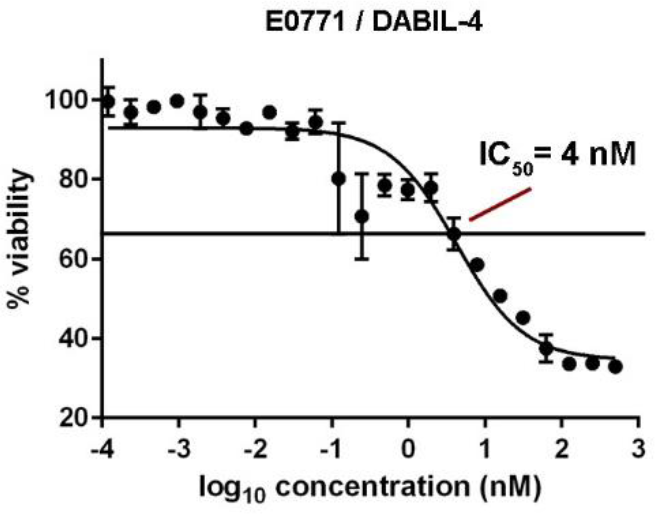
DABIL-4 shows cytotoxic activity against IL-4R^+^ E0771 TNBC cell line. Activity of DABIL-4 in MTS assay showed an IC_50_ of 4 nM.

**Fig S3.**
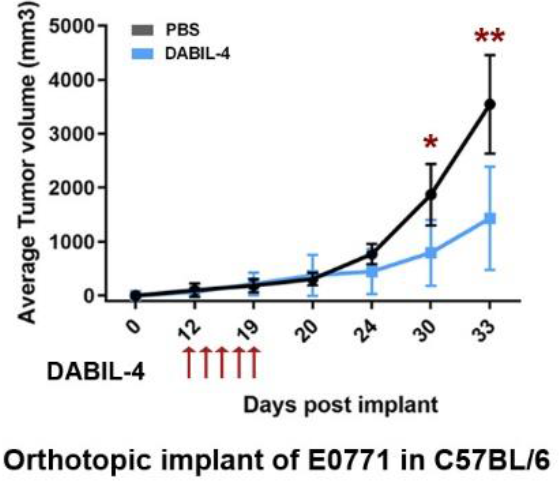
DABIL-4 exhibits anti-tumor activity in E0771 adenocarcinoma model in C57BL/6 mice. 50,000 E0771 cells were orthotopically implanted in C57BL/6 mice. Starting day 11, DABIL-4 was given i.p. on alternate days for total of 5 doses. Tumor volumes were measured using electronic Vernier Calipers. Statistical significance was calculated by two-tailed unpaired student t-test considering an unequal distribution. Data are shown as mean ± SD. *P < 0.05, **P < 0.01.

**Fig S4.**
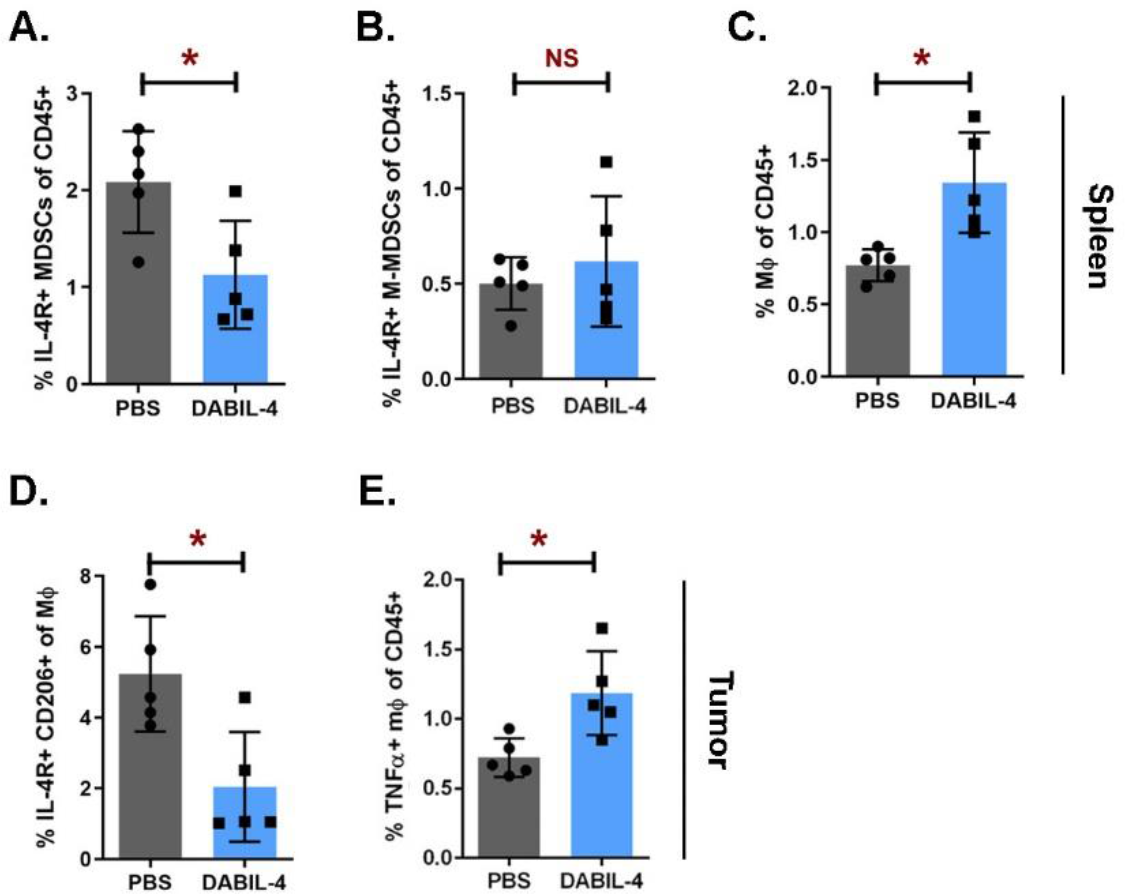
DABIL-4 treatment modulates myeloid cell populations in spleen and tumor microenvironment. Single cell suspensions of spleen and tumors were stained and analyzed by flow cytometry (n=5). We evaluated differences in the population of (A) IL-4R^+^ MDSCs of CD45^+^, (B) IL-4R^+^ M-MDSCs of CD45^+^, (C) macrophages of CD45^+^. (D) IL-4R^+^ CD206^+^ expression on macrophages and, (E) TNFα^+^ macrophages of CD45^+^. All panels correspond to day 25 post tumor implantation. Statistical significance between the groups was assessed by two-tailed unpaired student t-test considering an unequal distribution. Data are represented as mean ± SD. *P < 0.05, NS=non-significant.

**Fig S5.**
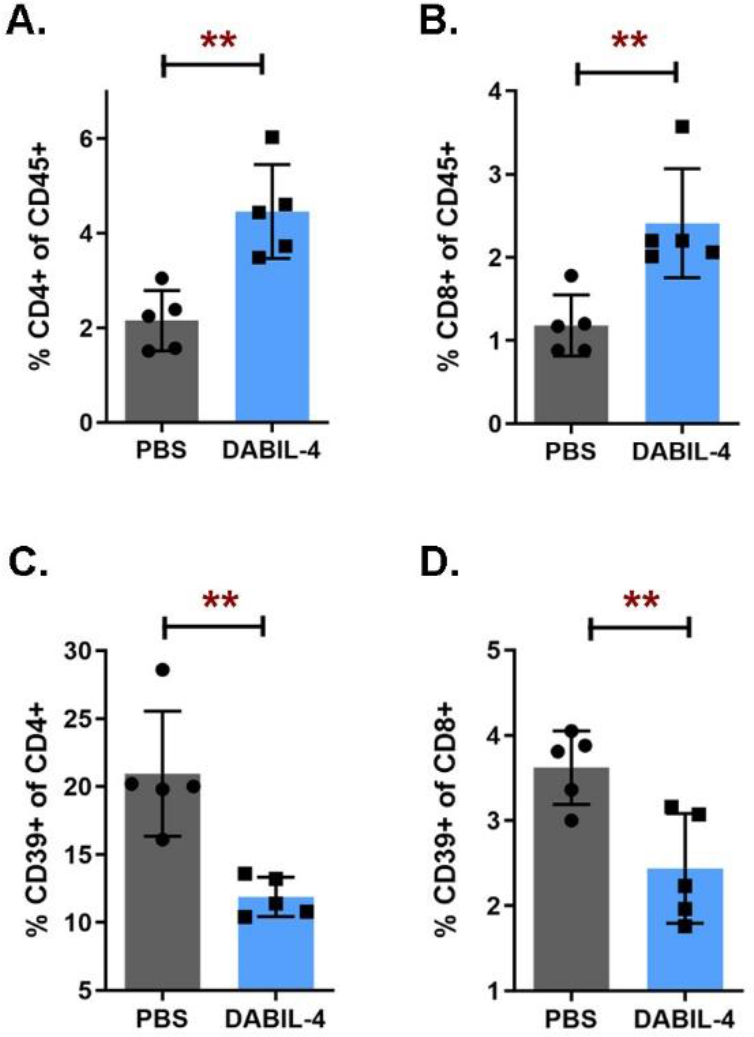
DABIL-4 administration promotes accumulation of lymphoid cell populations in spleen. Single cell suspensions of spleen were stained and analyzed by flow cytometry (n=5). We evaluated differences in the population of (A) CD4^+^ and (B) CD8^+^ T-cells of CD45^+^ cells. We also noted differences in (C) CD39 expression upon CD4^+^ and (D) CD8^+^ T-cells. All panels correspond to day 25 post tumor implantation. Statistical significance between the groups was assessed by two-tailed unpaired student t-test considering an unequal distribution. Data are represented as mean ± SD. *P < 0.05, **P < 0.01.

**Fig S6.**
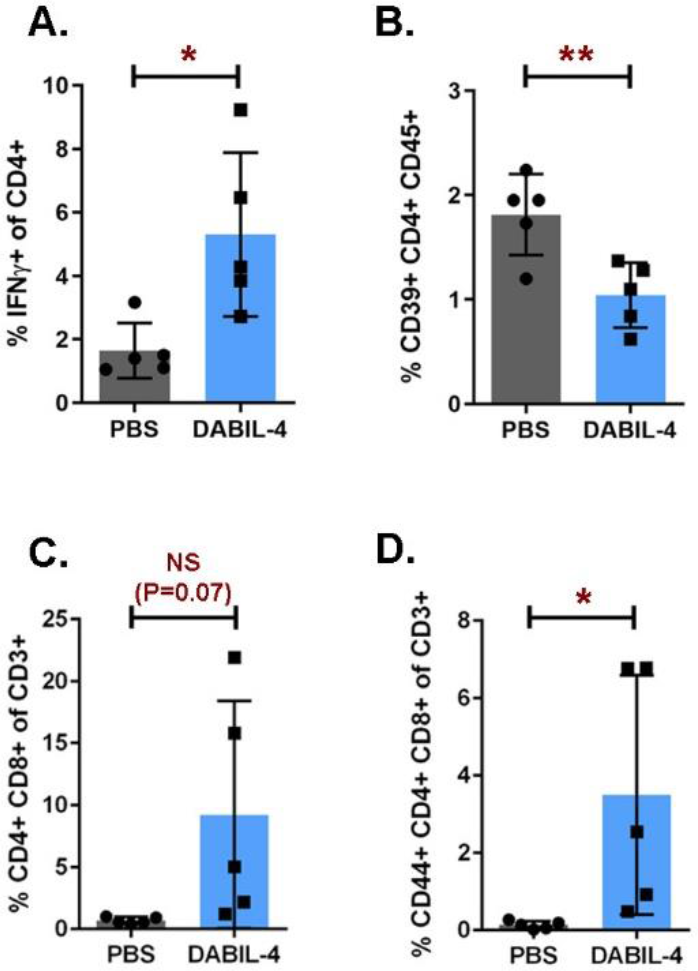
DABIL-4 promotes accumulation of activated effector cell population in tumor microenvironment. Single cell suspensions of tumors were stained and analyzed by flow cytometry (n=5). We evaluated differences in the population of **(A)** IFNγ production by CD4^+^, (B) CD39^+^ CD4^+^ of CD45^+^, (C) double-positive (DP) CD4^+^ CD8^+^ of CD3^+^ and (D) CD44^+^ DP of all lymphocytes. All panels except panel A correspond to day 17 post tumor implantation while panel A corresponds to day 25. Statistical significance between the groups was assessed by two-tailed unpaired student t-test considering an unequal distribution. Data are represented as mean ± SD. *P < 0.05, NS=non-significant.

## BIBILIOGRAPHY

1. S. Setrerrahmane, H. Xu, Tumor-related interleukins: old validated targets for new anti-cancer drug development. Mol Cancer 16, 153 (2017).

2. A. Suzuki, P. Leland, B. H. Joshi, R. K. Puri, Targeting of IL-4 and IL-13 receptors for cancer therapy. Cytokine 75, 79–88 (2015).

3. K. V. Bankaitis, B. Fingleton, Targeting IL4/IL4R for the treatment of epithelial cancer metastasis. Clin Exp Metastasis 32, 847–856 (2015).

4. M. Gaggianesi et al., IL4 Primes the Dynamics of Breast Cancer Progression via DUSP4 Inhibition. Cancer research 77, 3268–3279 (2017).

5. A. Konig et al., Determination of Interleukin-4, −5, −6, −8 and −13 in Serum of Patients with Breast Cancer Before Treatment and its Correlation to Circulating Tumor Cells. Anticancer Res 36, 3123–3130 (2016).

6. K. T. Venmar, K. J. Carter, D. G. Hwang, E. A. Dozier, B. Fingleton, IL4 receptor ILR4alpha regulates metastatic colonization by mammary tumors through multiple signaling pathways. Cancer research 74, 4329–4340 (2014).

7. K. T. Venmar, D. W. Kimmel, D. E. Cliffel, B. Fingleton, IL4 receptor alpha mediates enhanced glucose and glutamine metabolism to support breast cancer growth. Biochim Biophys Acta 1853, 1219–1228 (2015).

8. L. A. Mina, S. Lim, S. W. Bahadur, A. T. Firoz, Immunotherapy for the Treatment of Breast Cancer: Emerging New Data. Breast Cancer (Dove Med Press) 11, 321–328 (2019).

9. T. G. Lyons, Targeted Therapies for Triple-Negative Breast Cancer. Curr Treat Options Oncol 20, 82 (2019).

10. T. G. Lyons, T. A. Traina, Emerging Novel Therapeutics in Triple-Negative Breast Cancer. Adv Exp Med Biol 1152, 377–399 (2019).

11. L. S. Cheung et al., Second-generation IL-2 receptor-targeted diphtheria fusion toxin exhibits antitumor activity and synergy with anti-PD-1 in melanoma. Proceedings of the National Academy of Sciences of the United States of America 116, 3100–3105 (2019).

12. C. N. Johnstone et al., Functional and molecular characterisation of EO771.LMB tumours, a new C57BL/6-mouse-derived model of spontaneously metastatic mammary cancer. Dis Model Mech 8, 237–251 (2015).

13. M. Yamaizumi, E. Mekada, T. Uchida, Y. Okada, One molecule of diphtheria toxin fragment A introduced into a cell can kill the cell. Cell 15, 245–250 (1978).

14. B. A. Pulaski, S. Ostrand-Rosenberg, Mouse 4T1 breast tumor model. Curr Protoc Immunol Chapter 20, Unit 20.22 (2001).

15. G. Pascual et al., Targeting metastasis-initiating cells through the fatty acid receptor CD36. Nature 541, 41–45 (2017).

16. T. Suzuki et al., Early growth responsive gene 3 in human breast carcinoma: a regulator of estrogen-meditated invasion and a potent prognostic factor. Endocr Relat Cancer 14, 279–292 (2007).

17. R. Pio, Z. Jia, V. T. Baron, D. Mercola, Early growth response 3 (Egr3) is highly over-expressed in non-relapsing prostate cancer but not in relapsing prostate cancer. PloS one 8, e54096 (2013).

18. M. H. Chien et al., KSRP suppresses cell invasion and metastasis through miR-23a-mediated EGR3 mRNA degradation in non-small cell lung cancer. Biochim Biophys Acta Gene Regul Mech 1860, 1013–1024 (2017).

19. M. Pavlovic et al., Enhanced MAF Oncogene Expression and Breast Cancer Bone Metastasis. J Natl Cancer Inst 107, djv256 (2015).

20. M. Yao et al., CCR2 Chemokine Receptors Enhance Growth and Cell-Cycle Progression of Breast Cancer Cells through SRC and PKC Activation. Mol Cancer Res 17, 604–617 (2019).

21. S. T. Boyle, J. W. Faulkner, S. R. McColl, M. Kochetkova, The chemokine receptor CCR6 facilitates the onset of mammary neoplasia in the MMTV-PyMT mouse model via recruitment of tumor-promoting macrophages. Mol Cancer 14, 115 (2015).

22. N. Erin et al., Bidirectional effect of CD200 on breast cancer development and metastasis, with ultimate outcome determined by tumor aggressiveness and a cancer-induced inflammatory response. Oncogene 34, 3860–3870 (2015).

23. F. Talebian et al., Melanoma cell expression of CD200 inhibits tumor formation and lung metastasis via inhibition of myeloid cell functions. PloS one 7, e31442 (2012).

24. S. A. DuPre, K. W. Hunter, Jr., Murine mammary carcinoma 4T1 induces a leukemoid reaction with splenomegaly: association with tumor-derived growth factors. Exp Mol Pathol 82, 12–24 (2007).

25. F. Veglia, M. Perego, D. Gabrilovich, Myeloid-derived suppressor cells coming of age. Nature immunology 19, 108–119 (2018).

26. K. Nakamura, M. J. Smyth, Myeloid immunosuppression and immune checkpoints in the tumor microenvironment. Cell Mol Immunol 17, 1–12 (2020).

27. M. Dysthe, R. Parihar, Myeloid-Derived Suppressor Cells in the Tumor Microenvironment. Adv Exp Med Biol 1224, 117–140 (2020).

28. H. W. Wang, J. A. Joyce, Alternative activation of tumor-associated macrophages by IL-4: priming for protumoral functions. Cell Cycle 9, 4824–4835 (2010).

29. S. Ostrand-Rosenberg, P. Sinha, Myeloid-derived suppressor cells: linking inflammation and cancer. Journal of immunology (Baltimore, Md.: 1950) 182, 4499–4506 (2009).

30. I. Perrot et al., Blocking Antibodies Targeting the CD39/CD73 Immunosuppressive Pathway Unleash Immune Responses in Combination Cancer Therapies. Cell Rep 27, 2411–2425.e2419 (2019).

31. J. Desfrancois et al., Increased frequency of nonconventional double positive CD4CD8 alphabeta T cells in human breast pleural effusions. International journal of cancer 125, 374–380 (2009).

32. M. L. Clenet, F. Gagnon, A. C. Moratalla, E. C. Viel, N. Arbour, Peripheral human CD4(+)CD8(+) T lymphocytes exhibit a memory phenotype and enhanced responses to IL-2, IL-7 and IL-15. Sci Rep 7, 11612 (2017).

33. F. Lakkis et al., Interleukin 4 receptor targeted cytotoxicity: genetic construction and in vivo immunosuppressive activity of a diphtheria toxin-related murine interleukin 4 fusion protein. European journal of immunology 21, 2253–2258 (1991).

34. P. Leland et al., Human breast carcinoma cells express type II IL-4 receptors and are sensitive to antitumor activity of a chimeric IL-4-Pseudomonas exotoxin fusion protein in vitro and in vivo. Mol Med 6, 165–178 (2000).

35. V. Kumar, S. Patel, E. Tcyganov, D. I. Gabrilovich, The Nature of Myeloid-Derived Suppressor Cells in the Tumor Microenvironment. Trends in immunology 37, 208–220 (2016).

36. I. H. Younos, A. J. Dafferner, D. Gulen, H. C. Britton, J. E. Talmadge, Tumor regulation of myeloid-derived suppressor cell proliferation and trafficking. Int Immunopharmacol 13, 245–256 (2012).

37. J. R. Warner, The economics of ribosome biosynthesis in yeast. Trends Biochem Sci 24, 437–440 (1999).

38. V. Bronte et al., Recommendations for myeloid-derived suppressor cell nomenclature and characterization standards. Nat Commun 7, 12150 (2016).

39. L. Yang et al., Abrogation of TGF beta signaling in mammary carcinomas recruits Gr-1+CD11b+ myeloid cells that promote metastasis. Cancer Cell 13, 23–35 (2008).

40. I. Marigo et al., Tumor-induced tolerance and immune suppression depend on the C/EBPbeta transcription factor. Immunity 32, 790–802 (2010).

41. S. Mabuchi et al., Uterine Cervical Cancer Displaying Tumor-Related Leukocytosis: A Distinct Clinical Entity With Radioresistant Feature. JNCI: Journal of the National Cancer Institute 106 (2014).

42. D. Di Mitri, A. Toso, A. Alimonti, Molecular Pathways: Targeting Tumor-Infiltrating Myeloid-Derived Suppressor Cells for Cancer Therapy. Clin Cancer Res 21, 3108–3112 (2015).

43. A. Inoue, Y. Omoto, Y. Yamaguchi, R. Kiyama, S. I. Hayashi, Transcription factor EGR3 is involved in the estrogen-signaling pathway in breast cancer cells. J Mol Endocrinol 32, 649–661 (2004).

44. A. Bisgin, W. J. Meng, G. Adell, X. F. Sun, Interaction of CD200 Overexpression on Tumor Cells with CD200R1 Overexpression on Stromal Cells: An Escape from the Host Immune Response in Rectal Cancer Patients. J Oncol 2019, 5689464 (2019).

45. S. Shresta, P. Goda, R. Wesselschmidt, T. J. Ley, Residual cytotoxicity and granzyme K expression in granzyme A-deficient cytotoxic lymphocytes. The Journal of biological chemistry 272, 20236–20244 (1997).

46. K. Asadullah, W. Sterry, H. D. Volk, Interleukin-10 therapy--review of a new approach. Pharmacol Rev 55, 241–269 (2003).

