## Supplemental texts and Figures for "IL-4 receptor targeting as an effective immunotherapy against triple-negative breast cancer"

**This PDF file includes:**

Supplementary text (Materials and Methods)

Figures legends for S1 to S6

SI Bibliography

Figures S1 to S6

### MATERIALS AND METHODS

#### Plasmids, Bacterial Strains, and Cloning

*E. coli-C. diphtheriae* shuttle vector pKN2.6Z LC127 was constructed in the lab (1). Murine IL-4 sequence was codon optimized as per *C. diphtheriae* codon usage table and was synthesized by GenScript. hIL-2 sequence in pKN2.6z LC127 with mIL-4 sequence to generate pKN2.6z LC128 using Gibson assembly (NEB). Primers were specifically designed to introduce TEV (Tobacco Etch Virus)-protease sequence between murine IL-4 and His-6 sequence. The construct was then transformed into either chemically competent *Escherichia coli* DH5α strain (NEB) or electrocompetent *C. diphtheriae* C7s(−)*tox*-.

#### Fermenter Protein purification

*C. diphtheriae* non-lysogenic non-toxigenic C7s(-)*tox*- strain transformed with pKN2.6Z-LC128 was grown in CY medium in a fermenter (Bioflo/Celligen 110) as previously described (1). At OD ~12-15, culture was harvested, supernatant was collected and concentrated 20-fold by tangential flow filtration and a 30kD hollow fiber membrane (Spectrum). The flow concentrate was then diafiltered against 2 L of 50 mM $NaH_2PO_4$, 500 mM NaCl, 50 mM imidazole (pH 7.4). Concentrate was then adsorbed on to a HisTrap HP column (GE Healthcare) and the protein was eluted with 50 mM $NaH_2PO_4$, 500 mM NaCl, 500 mM imidazole (pH 7.4). The eluate was again concentrated using a 10 kDa Amicon Ultra-15 centrifugal units (Millipore Sigma) and separated over a HiPrep 26/60 Sephacryl S100-HR sizing column (GE Healthcare Life Sciences). The protein was eluted with PBS in 5 mL fractions and analyzed by SDS-PAGE. The protein concentration was then estimated and aliquots were stored at -80°C in PBS.

#### Cell lines, media and growth conditions

Murine origin triple-negative breast cancer cell lines 4T1 (laboratory stock) and EO771 (purchased from CH3 biosystems) were grown in DMEM F12 (Corning) and RPMI-1640 (Corning) + 0.1% HEPES respectively supplemented with 10% heat-inactivated FBS (Gibco) and 1% antibacterial antimycotic solution (Sigma Aldrich). The cell lines were grown in a humidified incubator at 37°C under 5% $CO_2$ atmosphere.

#### MTS cell viability Assay

For MTS assay, 5000 4T1 cells were plated per well in triplicate in 200 ul volume in 96-well plates, allowed to adhere and treated with 2-fold serial dilutions of DABIL-4 fusion toxin or no drug. After 48 h incubation with the drug, [3-(4,5-dimethylthiazol-2-yl)-5-(3-carboxymethoxyphenyl)-2-(4-sulfophenyl)-2H-tetrazolium, inner salt; MTS reagent (Promega) was added to the individual wells. After 90-120 minutes, absorbance at 490 nM was recorded with the iMark Microplate Reader (Bio-Rad) to score cell viability.

#### Trypan blue dye exclusion assay

5000 4T1 cells/well were plated in triplicate in 1 ml volume in 12 well plates, treated with 4-fold serial dilutions of DABIL-4 fusion toxin or no drug, and incubated for 48 h. Cells were harvested, resuspended in trypan blue dye and counted using hemocytometer (2).

#### Western blots

For analyzing the protein samples, media supernatant/purified protein was boiled in 4x SDS loading buffer (Bio-Rad) and separated on pre-casted 4-15% SDS-PAGE gel (Bio-Rad). The separated proteins were transferred to activated PVDF membrane and probed with appropriate primary antibodies followed by secondary antibodies and detected by ECL reagent (Thermo fisher). Antibodies for western blots were purchased from different sources: anti-IL-2 (CST, Cat#12239), anti-His (Abcam, Cat# ab18184), anti-DT (Abcam, Cat# ab53828), anti-total PARP (CST, Cat#9532), anti-cleaved PARP (CST, Cat#5625), anti-BCL-xL (CST, Cat#), anti-cleaved caspase-3 (CST, Cat#), anti-β-actin (Sigma, Cat# A5441). CST stands for Cell signaling technology (Danvers, MA, USA).

**Caspase activity assay**

4T1 cells were seeded at the density of 1000 cells/well. Post adherence, cells were treated with no drug, 20 nM DABIL-4 and 2.5 µM Doxurubicin (as a positive control). Caspase activities in cells were quantified post 48 hours of treatment using Caspase-Glo® 3/7 Assay System (Promega) as per manufacturer's instructions.

For testing the expression of apoptosis associated markers (such as PARP, caspase-3 and BCL-xl), 4T1 cells were seeded in 6-well plates at a density of 5000 cells/well and treated with DABIL-4 or no drug. After 48 hours, cells were trypsinised, washed with PBS, and lysed with RIPA buffer (Thermo Fisher) for 30 min on ice. The lysates were then centrifuged at 13,000 rpm for 10 min at 4°C. The protein concentration was estimated as per Bradford assay and equal samples were loaded on SDS-PAGE gel (40 µg protein per lane). Western blot was performed as mentioned in the earlier section.

***In vivo* mouse experiments**

All animal experimental procedures were reviewed and approved by Johns Hopkins University Animal Care and Use Committee. For tumor generation, 10,000 4T1 cells in 0.1 mL Matrigel (50 % v/v in PBS; Corning) were orthotopically implanted into $8^{th}$ mammary fat pad of 6-week-old female Balb/c mice (Charles River Laboratories). At day 7, when tumor became palpable, PBS and DABIL-4

(10 µg/mice in 100 µL PBS) were administered via intraperitoneal route on alternate days thrice a week till the completion of the study. Tumor growth was assessed twice a week (Mon and Thurs) by measuring tumor volume using electronic caliper. Tumor volume was calculated suing the following equation = length X width X width X 0.5. Mice were sacrificed at the indicated time points. Tumor and spleen were isolated and weighed.

For single cell suspension preparation, tumors were minced and digested with collagenase D and DNase I at 37 C for 1 h while spleens were passed through 100 µm mesh filters. Red blood cells were lysed using RBC lysis buffer (Biolegend).

**Flow cytometry**

Single-cell suspensions from the tumor and spleen were stained with trypan blue and manually scored for viability. 1 million cells were incubated with purified anti-mouse CD16/32 antibody (Biolegend, Cat# 101320) in FACS buffer (eBiosciences, Cat# 422226) and stained with two panels of antibodies (Biolegend unless otherwise indicated); (1) Lymphoid panel: APC/Cy7 CD45, BV785 CD3, BUV563 CD4 (BD), AF700 CD8a, BV711 CD25, PE-Texas Red CD39, BV650 CD44, BV421 PD-L1, (2) Myeloid Panel: APC/Cy7 CD45, BV650 CD11b, BV605 Ly6G (BD), PE/CY7 Ly6C, BV711 CD115, PE CD124, BUV396 IA/IE (BD), APC F4/80, FITC CD86,. Zombie-Aqua fixable viability dye was included in both the panels to select for viable cells. Post surface staining, cells were fixed in fixation buffer and intracellular staining was performed using transcription factor buffer set followed by staining with FITC Foxp3 (lymphoid panel) and PerCP/Cy5.5 CD206 (myeloid panel). The stained samples were acquired on LSRFortessa™ X-20 Cell Analyzer (BD) and data were analyzed using FlowJo v.10 software (Tree Star). To exclude debris and aggregates, FSC-A versus SSC-A gate and to exclude duplets, FSC-W and SSC-H gates were designed. Inside this latter gate, dead cells were excluded using the Zombie-red fixable viability dye. All reagents were from Biolegend unless otherwise indicated.

**T-cell stimulation and Intracellular Cytokine staining**

For *in vitro* stimulations, harvested cells were incubated with cell activation cocktail (PMA-Ionomycin without Brefeldin) and monesin in RPMI-1640 media supplemented with 10% FBS at 37 C for 4 hours. Post stimulation, cells were surface stained with APC/Cy7 CD45, BV785 CD3, BUV563 CD4, AF700 CD8a as described earlier. Intracellular cytokine staining was performed with cyto-fast™ fix/perm buffer set as per the manufacturer's protocol and labeled with BV421 IFNϒ. Samples were acquired on LSRFortessa™ X-20 Cell Analyzer (BD) and data was analyzed using FlowJo v.10 software (Tree Star). Zombie aqua fixable dye was included to score viability. All reagents were supplied by Biolegend unless otherwise indicated.

**Lung metastasis assay**

To quantify metastases, lungs were isolated from tumor bearing mice, excised, minced and digested with collagenase D and DNase I to prepare a single cell suspension. The suspension was plated in 6-well plates in DMEM F-12 media supplemented with s-guanidine and incubated in a humidified incubator at 37 under 5% CO2 for 10-14 days. Following selection, colonies were formalin-fixed, stained with crystal violet prior to manual counting. Plates were also photographed for record keeping.

**RNA isolation and NanoString analysis**

RNA was isolated from 3 4T1 tumors from both DABIL-4 treated and untreated group by using RNeasy Mini Kit (Qiagen) as per manufacturer's protocol. All RNA samples passed quality control (assessed by OD 260/280) and were analyzed by nCounter murine PanCancer Immune Profiling

Panel according to manufacturer's protocol at Johns Hopkins Transcriptomics and Deep Sequencing Core Facility (NanoString Technologies). Raw data was normalized using the nSolver 3.0 analysis software (NanoString Technologies). Gene expression (represented in $log_2$) was calculated as the mean and data was imported to GraphPad Prism software for statistical analysis and graphical representation.

#### SUPPLEMENTARY BIBILOGRAPHY


1. L. S. Cheung *et al.*, Second-generation IL-2 receptor-targeted diphtheria fusion toxin exhibits antitumor activity and synergy with anti-PD-1 in melanoma. *Proceedings of the National Academy of Sciences of the United States of America* **116**, 3100-3105 (2019).
2. W. Strober, Trypan blue exclusion test of cell viability. *Curr Protoc Immunol* **Appendix 3**, Appendix 3B (2001).

### SUPPLEMENTARY FIGURES

#### Fig. S1

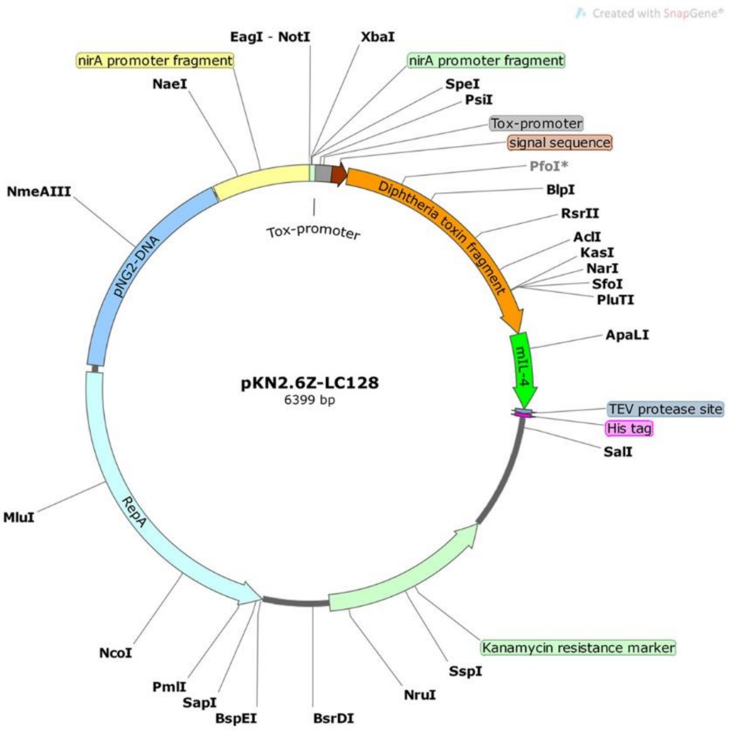


**Fig S1.** pKN2.6Z-LC128 shuttle vector plasmid map.

Fig. S2

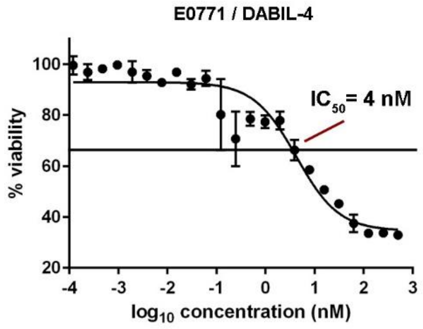


**Fig S2. DABIL-4 shows cytotoxic activity against IL-4R$^{+}$ E0771 TNBC cell line.** Activity of DABIL-4 in MTS assay showed an $IC_{50}$ of 4 nM.

### Fig. S3

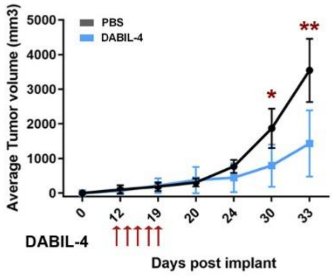


Orthotopic implant of E0771 in C57BL/6

**Fig S3. DABIL-4 exhibits anti-tumor activity in E0771 adenocarcinoma model in C57BL/6 mice.** 50,000 E0771 cells were orthotopically implanted in C57BL/6 mice. Starting day 11, DABIL-4 was given i.p. on alternate days for total of 5 doses. Tumor volumes were measured using electronic Vernier Calipers. Statistical significance was calculated by two-tailed unpaired student t-test considering an unequal distribution. Data are shown as mean ± SD. *P < 0.05, **P < 0.01.

**Fig. S4**

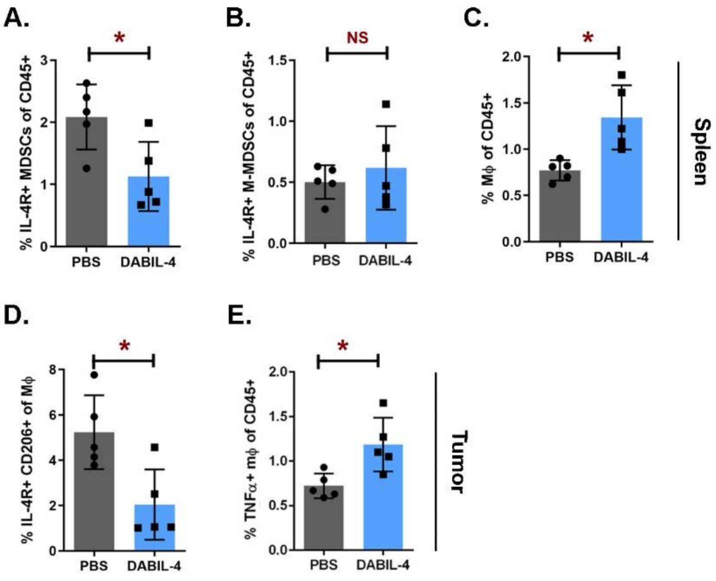


**Fig S4. DABIL-4 treatment modulates myeloid cell populations in spleen and tumor microenvironment.** Single cell suspensions of spleen and tumors were stained and analyzed by flow cytometry (n=5). We evaluated differences in the population of **(A)** IL-4R$^+$ MDSCs of CD45$^+$, **(B)** IL-4R$^+$ M-MDSCs of CD45$^+$, (**C)** macrophages of CD45$^+$. **(D)** IL-4R$^+$ CD206$^+$ expression on macrophages and, **(E)** TNFα$^+$ macrophages of CD45$^+$. All panels correspond to day 25 post tumor implantation. Statistical significance between the groups was assessed by two-tailed unpaired student t-test considering an unequal distribution. Data are represented as mean ± SD. *P < 0.05, NS=non-significant.

**Fig. S5**

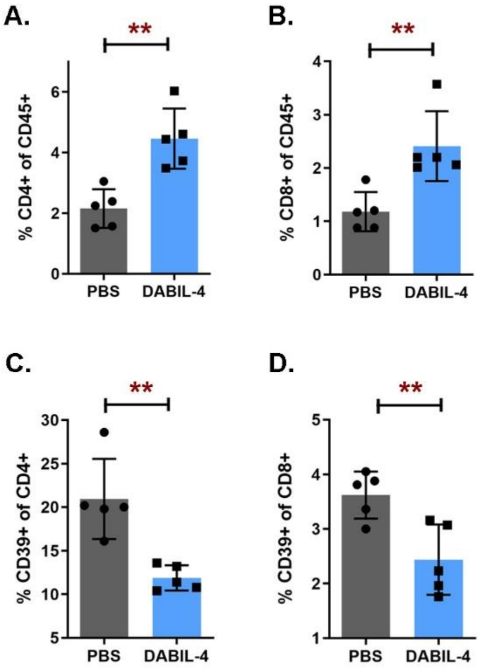


**Fig S5. DABIL-4 administration promotes accumulation of lymphoid cell populations in spleen.** Single cell suspensions of spleen were stained and analyzed by flow cytometry (n=5). We evaluated differences in the population of **(A)** $CD4^+$ and **(B)** $CD8^+$ T-cells of $CD45^+$ cells. We also noted differences in **(C)** CD39 expression upon $CD4^+$ and **(D)** $CD8^+$ T-cells. All panels correspond to day 25 post tumor implantation. Statistical significance between the groups was assessed by two-tailed unpaired student t-test considering an unequal distribution. Data are represented as mean ± SD. *P < 0.05, **P < 0.01.

#### Fig. S6

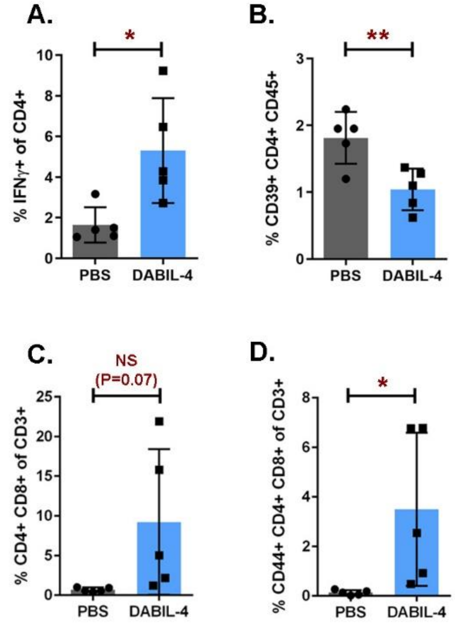


**Fig S6. DABIL-4 promotes accumulation of activated effector cell population in tumor microenvironment.** Single cell suspensions of tumors were stained and analyzed by flow cytometry (n=5). We evaluated differences in the population of **(A)** IFNγ production by $CD4^+$, (B) $CD39^+$ $CD4^+$ of $CD45^+$, (C) double-positive (DP) $CD4^+$ $CD8^+$ of $CD3^+$ and (D) $CD44^+$ DP of all lymphocytes. All panels except panel A correspond to day 17 post tumor implantation while panel A corresponds to day 25. Statistical significance between the groups was assessed by two-tailed unpaired student t-test considering an unequal distribution. Data are represented as mean ± SD. $*P < 0.05$, NS=non-significant.
